# Modeling Rotational Fluoroquinolone Therapy as a Novel Treatment for Ophthalmic MRSA Infections

**DOI:** 10.1101/2025.06.22.660934

**Authors:** Alejandro Storper, Darlene Miller, Xi Huo

## Abstract

Methicillin-resistant *Staphylococcus aureus* (MRSA) increasingly undermines the effectiveness of topical fluoroquinolone monotherapy in ophthalmic infections. We present a theoretical evaluation of rotational fluoroquinolone therapy, in which moxifloxacin and trovafloxacin are alternated to enhance bacterial suppression and mitigate resistance. Experimental time-kill data were fit using Bayesian parameter estimation via Markov Chain Monte Carlo (MCMC) to derive drug-specific growth and kill parameters, which were integrated into spatiotemporal pharmacodynamic models. These models incorporate radial anterior chamber geometry, intraocular diffusion, and aqueous humor pharmacokinetics governed by a circadian clearance function. Structural docking simulations of topoisomerase IV mutants reveal that common resistance mutations disrupt moxifloxacin binding while preserving trovafloxacin affinity, supporting a collateral-sensitivity mechanism. Under high-resistance conditions, rotational therapy improves bacterial clearance compared to monotherapy by increasing heterogeneity in antimicrobial exposure fronts and reducing overall bacterial burden. By alternating fluoroquinolones with complementary resistance profiles, rotational therapy leverages physiological clearance rhythms and spatial drug gradients to enhance anterior-segment efficacy, providing a quantitative foundation for its use as a rational alternative to monotherapy in ocular MRSA management.

## Introduction

Methicillin-resistant *Staphylococcus aureus* (MRSA) is a Gram-positive, facultative anaerobe and coccoid bacterium that causes both hospital- and community-acquired infections and can lead to severe ocular disease, including corneal ulcers and vision loss^1–5^. Postoperative procedures such as cataract surgery are associated with an increased risk of intraocular infection due to Gram-positive organisms, particularly in the absence of antibiotic prophylaxis^6^.

MRSA displays high rates of resistance to traditional staphylococcal antibiotics, particularly within healthcare environments^7,8^. Consequently, fluoroquinolones, which are broad-spectrum bactericidal agents effective against both gram-positive and gram-negative organisms, have seen increased use, notably as topical prophylaxis in ophthalmic surgery^9,10^.

Trovafloxacin, a fourth-generation fluoroquinolone originally developed for systemic use, exhibits potent activity against Gram-positive bacteria^11–15^ due to its high affinity for topoisomerase IV^16^(Figure 1). Its efficacy against MRSA has been linked to unique 2,4-difluorophenyl and azabicyclo[3.1.0]hexyl substitutions, which enhance enzyme binding and broaden its antimicrobial spectrum^17^. Although it was withdrawn from systemic markets because of rare but severe idiosyncratic hepatotoxicity, regulators removed trovafloxacin from use^18,19^. Its strong *in vitro* potency and the markedly reduced systemic exposure expected from topical ophthalmic administration render it a promising candidate for theoretical modeling and repurposing in ocular infections^20,21^.

**Figure 1.**
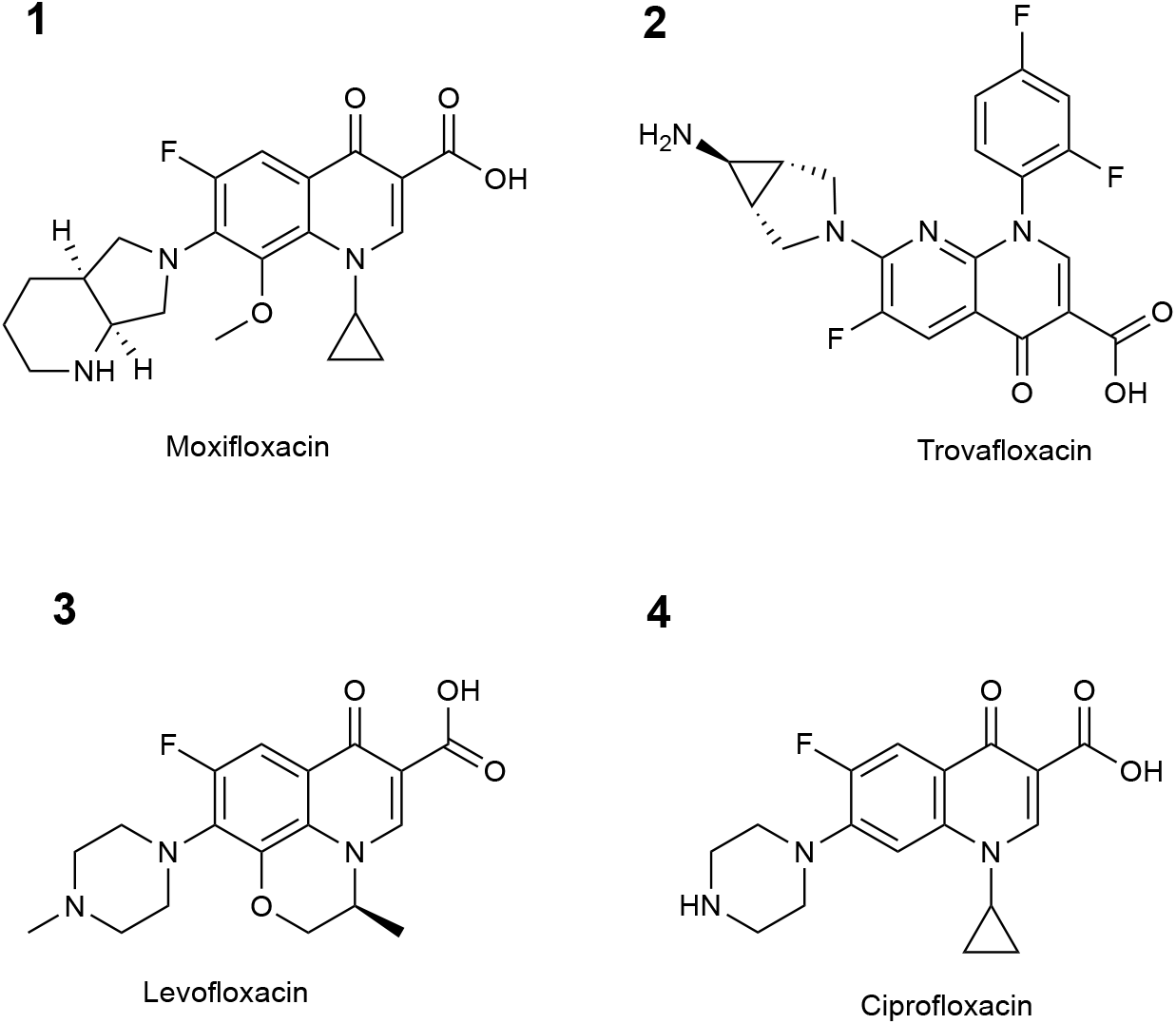
Chemical structures of (**1**) moxifloxacin, (**2**) trovafloxacin, (**3**) levofloxacin and (**4**) ciprofloxacin.

Moxifloxacin, a fourth-generation fluoroquinolone formulated for topical ophthalmic use, incorporates an 8-methoxy group and a bulky C-7 bicyclic side chain^22^ (Figure 1). This enhances lipophilicity, ocular tissue penetration, and topoisomerase IV binding, leading to higher, sustained concentrations in the cornea and aqueous humor^23^. Systemic absorption remains minimal, though QT-interval prolongation can occur at high plasma levels, particularly in patients with compromised aqueous outflow^24,25^.

Monotherapy with fluoroquinolones is common in ocular infections but can drive the emergence of resistant MRSA strains via selective pressure^26–29^. Resistance typically arises from mutations in quinolone-resistance determining regions (QRDRs) of *parC*, which decrease topoisomerase IV susceptibility^30–32^. Dual targeting strategies that combine inhibitors of DNA gyrase and topoisomerase IV have demonstrated enhanced bacterial efficacy^33,34^, although sustained monotherapy continues to select for resistant mutants.

Collateral sensitivity—where resistance to one antibiotic increases susceptibility to another—is a promising tactic to suppress resistance evolution^35–39^. For MRSA, mutations developed in response to moxifloxacin may reduce topoisomerase IV efficiency or stability, paradoxically increasing vulnerability to trovafloxacin, whose distinct structure may complement the altered binding pocket (Figure 2). Such vulnerabilities can be exploited via rotational therapy, imposing alternating selective pressures to hinder stable resistance establishment.

**Figure 2.**
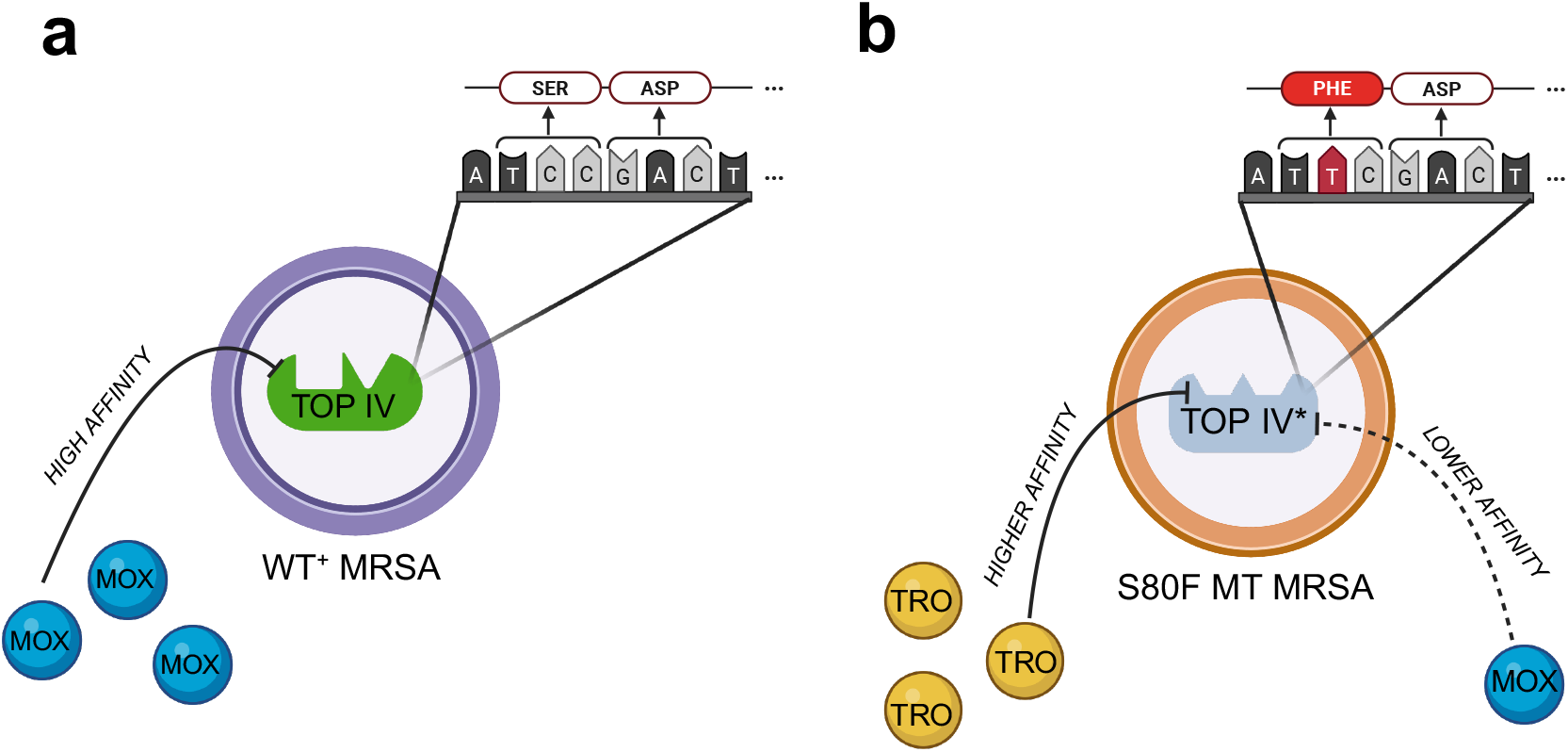
Collateral sensitivity induced by moxifloxacin resistance via *parC* S80F mutation in MRSA. (**a**) In wild-type (*parC*^+^) MRSA, moxifloxacin (MOX) binds topoisomerase IV (Top IV) with high affinity and inhibits DNA replication. (**b**) Selective pressure from moxifloxacin leads to a C→T point mutation in codon 80 of *parC* (TCC → TTC), resulting in an S80F amino acid substitution. This mutation reduces moxifloxacin binding to Top IV^*^, conferring resistance, while at the same time increasing trovafloxacin (TRO) affinity. Codon and amino acid sequences are shown for both wild-type and mutant strains.

Although rotational antibiotic therapy has been proposed in other infectious contexts to mitigate resistance evolution^40^, it has not been systematically explored for fluoroquinolones or MRSA. Our study builds upon the concept of collateral sensitivity and biochemical trade-offs associated with resistance mutations to evaluate whether alternating fluoroquinolone therapy can enhance bacterial suppression in ocular MRSA infections.

In this study, we use a mathematical model to test the hypothesis that rotational topical therapy between moxifloxacin and trovafloxacin, informed by circadian clearance kinetics and synergistic drug interactions, can more effectively suppress ocular MRSA compared to conventional monotherapy. We combine pharmacodynamic modeling, resistance evolution simulations, and molecular docking to evaluate the therapeutic potential of fluoroquinolone rotation in ophthalmic infections.

## Results

### Resistant MRSA isolates associated with baseline replication fitness cost

Baseline growth curves in the absence of antibiotic exhibited classical logistic dynamics for both isolates. Fitted growth parameters (Table 1) reveal that the resistant isolate’s intrinsic replication rate is reduced by approximately 21 % relative to the susceptible isolate, reflecting a clear fitness cost associated with resistance. Although this fitness cost slows proliferation under drug-free conditions, the resistant strain’s survival advantage emerges under moxifloxacin exposure.

**Table 1.**
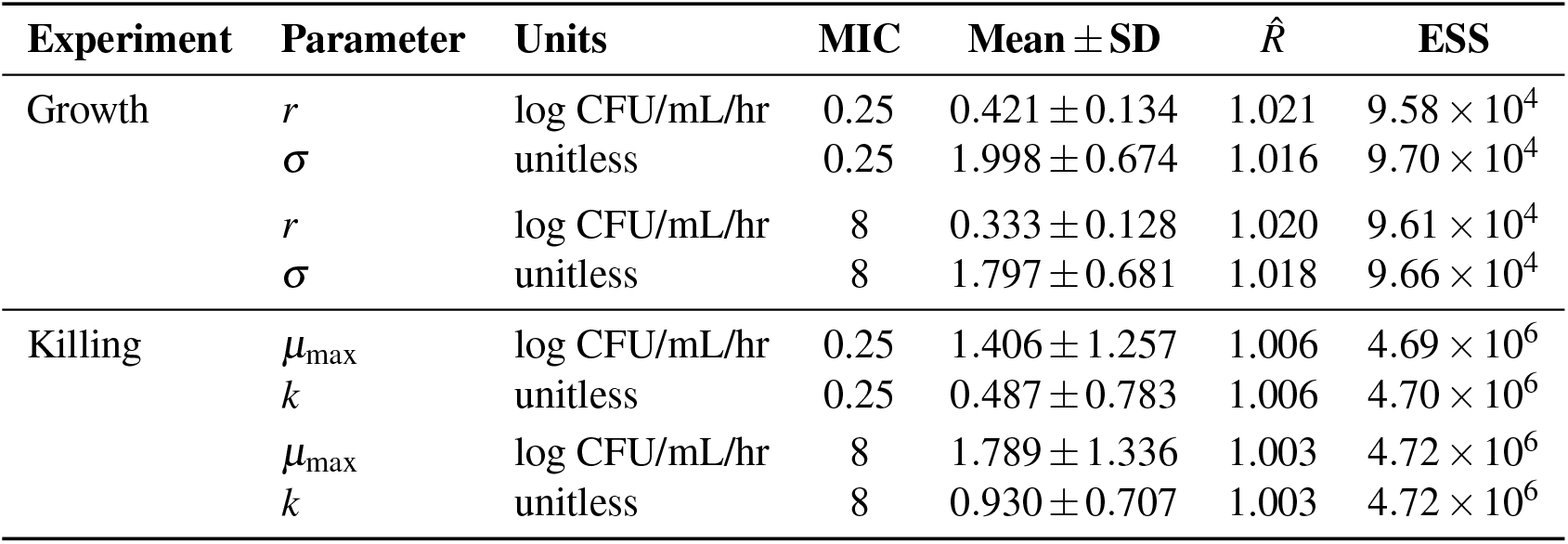
Summary of MCMC parameter estimates and convergence diagnostics.

| Experiment | Parameter | Units | MIC | Mean ± SD | $\hat{R}$ | ESS |
| --- | --- | --- | --- | --- | --- | --- |
| Growth | $r$ | log CFU/mL/hr | 0.25 | $0.421 \pm 0.134$ | 1.021 | $9.58 \times 10^4$ |
| | $\sigma$ | unitless | 0.25 | $1.998 \pm 0.674$ | 1.016 | $9.70 \times 10^4$ |
| | $r$ | log CFU/mL/hr | 8 | $0.333 \pm 0.128$ | 1.020 | $9.61 \times 10^4$ |
| | $\sigma$ | unitless | 8 | $1.797 \pm 0.681$ | 1.018 | $9.66 \times 10^4$ |
| Killing | $\mu_{\max}$ | log CFU/mL/hr | 0.25 | $1.406 \pm 1.257$ | 1.006 | $4.69 \times 10^6$ |
| | $k$ | unitless | 0.25 | $0.487 \pm 0.783$ | 1.006 | $4.70 \times 10^6$ |
| | $\mu_{\max}$ | log CFU/mL/hr | 8 | $1.789 \pm 1.336$ | 1.003 | $4.72 \times 10^6$ |
| | $k$ | unitless | 8 | $0.930 \pm 0.707$ | 1.003 | $4.72 \times 10^6$ |

In kill-curve assays, the susceptible isolate showed no significant inhibition at sub-MIC concentrations but was rapidly eradicated at twice the MIC and above (Figure 3). By contrast, the resistant isolate only exhibited bacteriostatic suppression at its MIC and required concentrations four-fold above the MIC to achieve complete bactericidal activity (Figure 4).

**Figure 3.**
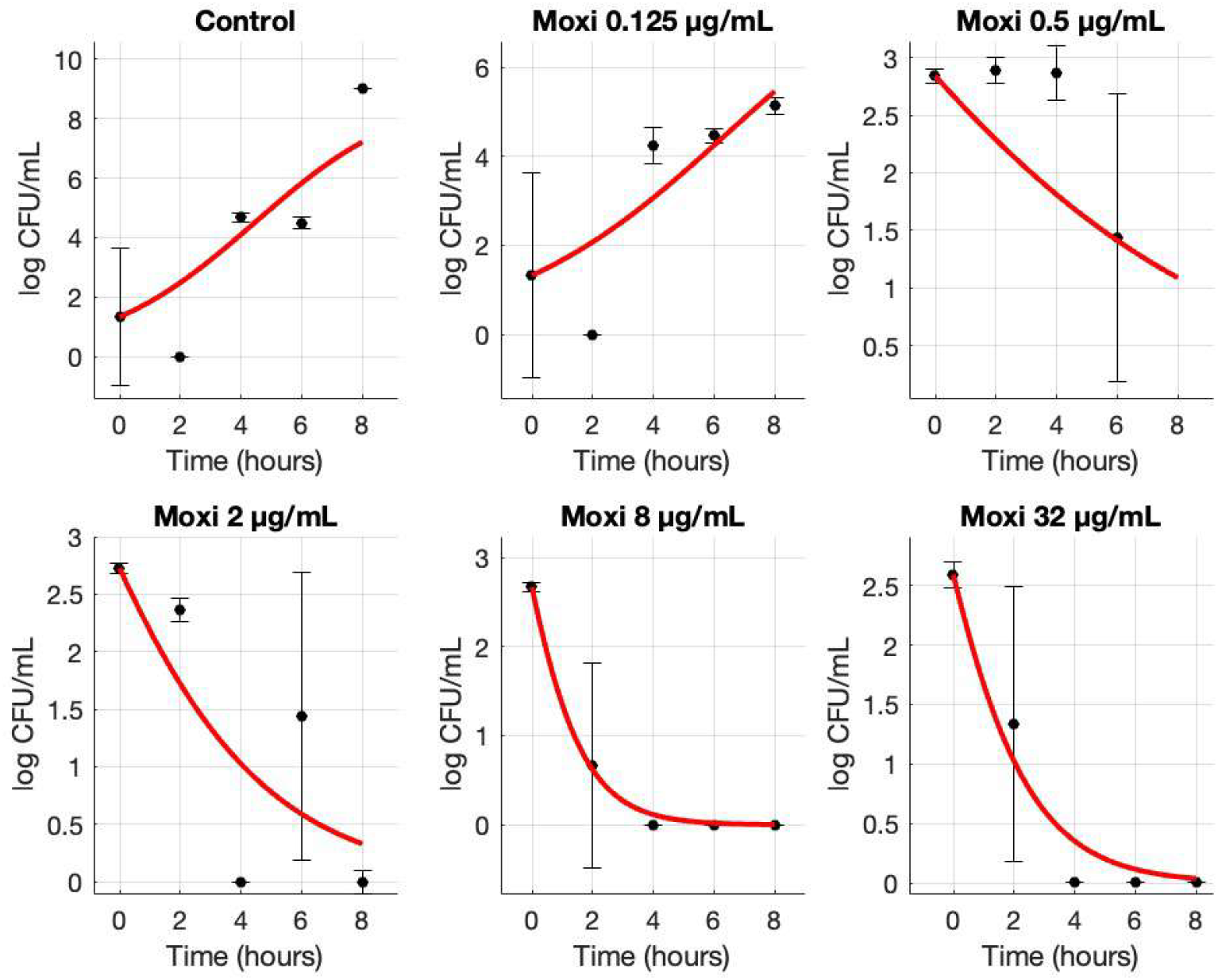
Growth and kill curves (mean ± SD) for a susceptible MRSA isolate upon treatment of varying moxifloxacin (Moxi) concentrations. CFU counts were conducted every 2 hours and were averaged among triplicates. Standard deviations are shown with error bars for each data point. The data points were then fit to their respective curve using MCMC.

**Figure 4.**
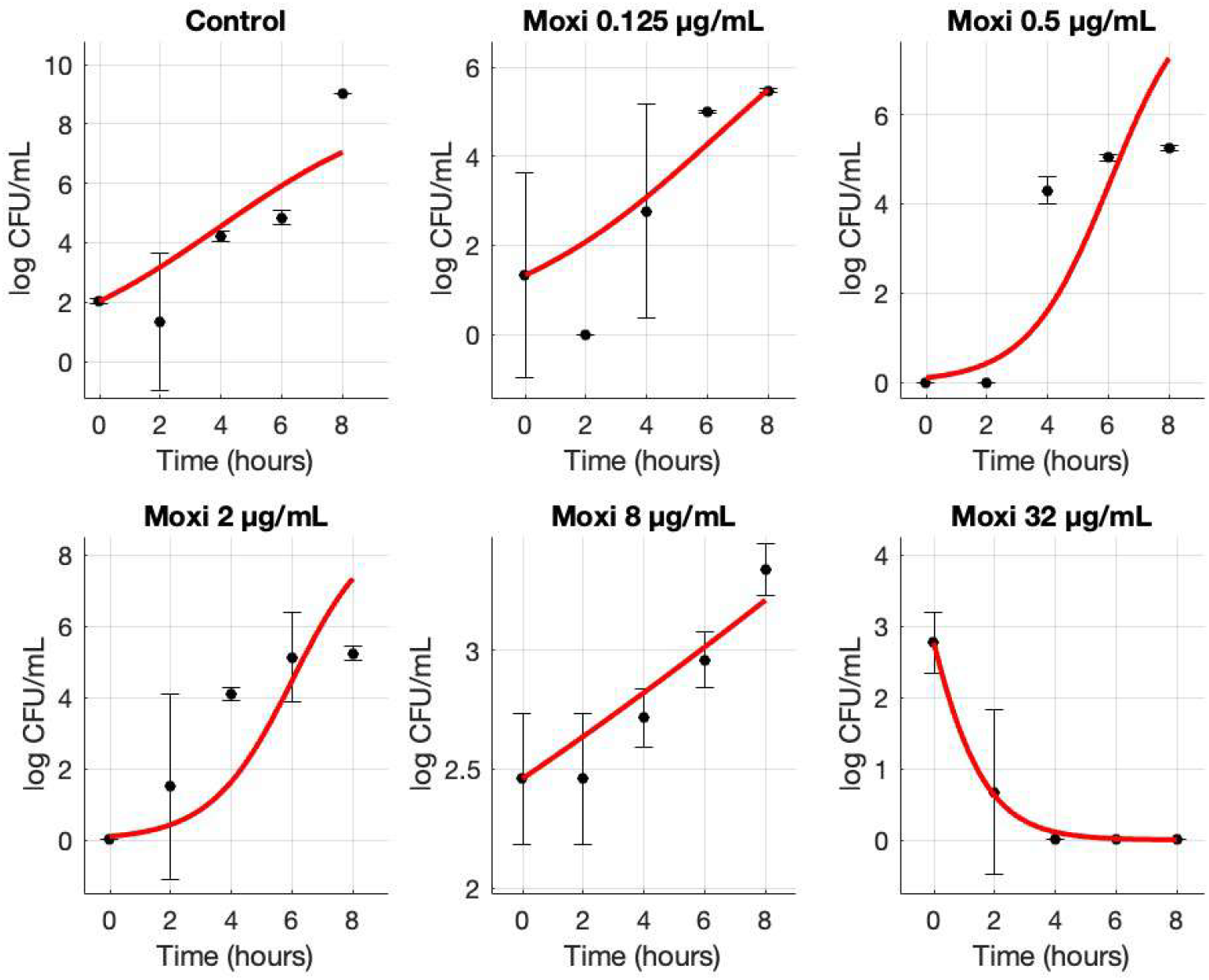
Growth and kill curves (mean ± SD) for a resistant MRSA isolate upon treatment of varying moxifloxacin (Moxi) concentrations. CFU counts were conducted every 2 hours and were averaged among triplicates. Standard deviations are shown with error bars for each data point. The data points were then fit to their respective curve using MCMC.

The concentration–kill rate relationship (Figure S1) demonstrates that the resistant isolate exhibits both a higher maximal kill rate and a larger Hill coefficient compared with the susceptible isolate. Its elevated MIC shifts the dose–response curve to the right, requiring higher drug concentrations to achieve full bactericidal activity and resulting in a broader, attenuated response profile. These findings emphasize the trade-off between replication fitness costs in antibiotic-free conditions and enhanced pharmacodynamic performance under antibiotic pressure.

### Moxifloxacin-induced topoisomerase IV mutations confer collateral sensitivity to trovafloxacin

Molecular docking simulations were performed to evaluate the binding interactions of fluoroquinolones with *S. aureus* topoisomerase IV, both in the wild-type enzyme and in a clinically relevant mutant (S80F). While we initially tested several fluoroquinolones already available in topical ophthalmic formulation (Figure 1), none demonstrated significant differential binding between wild-type and mutant structures. In contrast, trovafloxacin uniquely exhibited a marked change in affinity, indicating a robust collateral sensitivity effect.

Docking analysis indicates that the S80F mutation markedly compromises moxifloxacin binding: its calculated affinity decreases from –6.15 kcal/mol in the wild-type enzyme to –5.39 kcal/mol in the mutant, reflecting both loss of key chemical interactions and steric incompatibility. In contrast, trovafloxacin retains strong binding under both conditions (–5.89 kcal/mol wild-type; –6.00 kcal/mol S80F), consistent with its ability to accommodate structural rearrangements in the pocket (Figure 5). Structural analyses of the docked complexes reveal a clear mechanistic explanation for these differences. Moxifloxacin’s rigid C-7 diazabicyclic amine induces steric clashes with Phe80 in the mutated binding pocket, disrupting even its lone hydrophobic contact and abolishing key hydrogen bonds, which sharply reduces binding affinity. In contrast, trovafloxacin’s smaller, five-membered nitrogenous ring at C-7 engages directly with Phe80 through multiple hydrophobic and *π− π* stacking interactions, while its planar 2,4-difluorophenyl group at N-1 and adjacent hydrogen bonds to neighboring residues remain intact, enabling robust binding to the altered active site (Figure S9).

**Figure 5.**
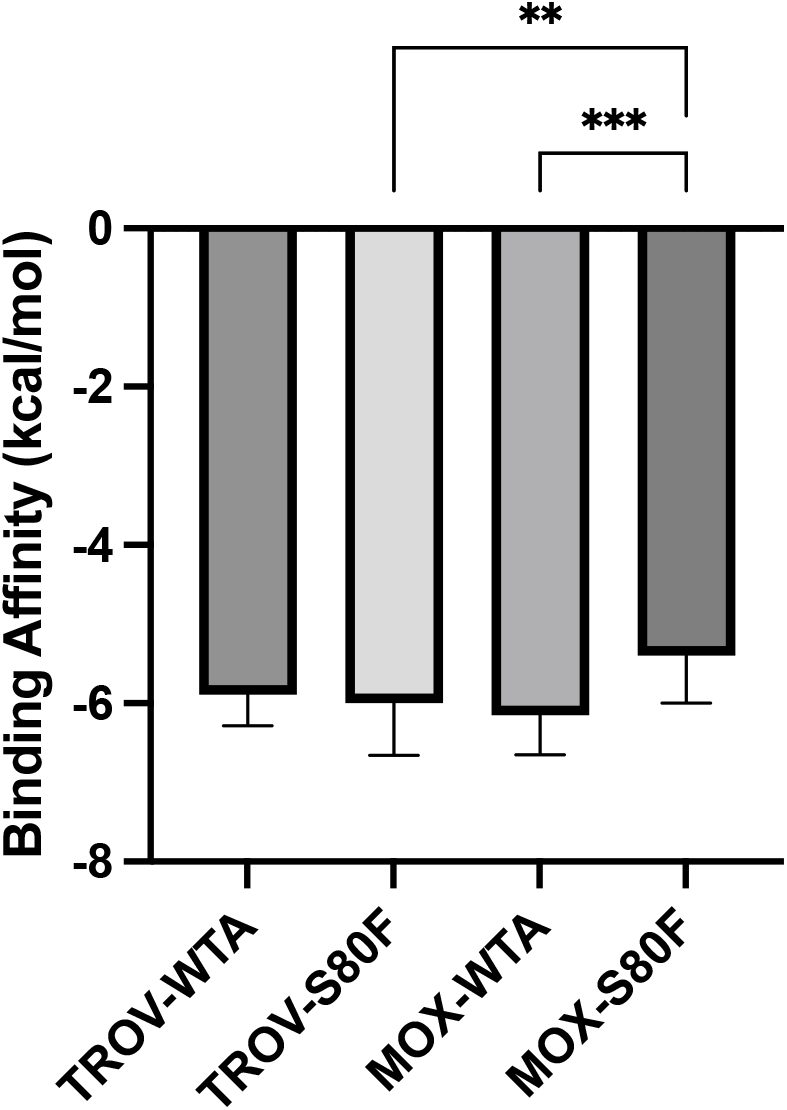
Binding affinities of moxifloxacin (MOX) and trovafloxacin (TROV) to topoisomerase IV subunit A in wild-type (WTA) and S80F-mutant forms. Both MOX and TROV exhibited strong binding to WTA of topoisomerase IV. The S80F mutation significantly reduced MOX binding affinity, while TROV binding remained relatively stable. Bars represent mean binding energies (kcal/mol) *±* standard deviation (s.d.), with *n* = 19–20 per group. One-way analysis of variance (ANOVA) was used to assess differences in group means under the assumption of normality and homogeneity of variances, confirmed by Brown–Forsythe and Bartlett’s tests (*P* > 0.05). Post hoc comparisons were conducted using Tukey’s honestly significant difference (HSD) test to correct for multiple comparisons. Asterisks denote adjusted *P* values: * *P* < 0.05, ** *P* < 0.01, *** *P* < 0.001. All tests were two-tailed and performed with *α* = 0.05.

These results demonstrate that the S80F mutation disproportionately disrupts moxifloxacin binding, while trovafloxacin retains robust affinity, suggesting a promising collateral sensitivity mechanism. This asymmetry provides a molecular rationale for using moxifloxacin to induce resistance mutations that inadvertently sensitize MRSA to trovafloxacin. Given that trovafloxacin is not currently formulated as an ophthalmic drop, our findings motivate further efforts to develop a topical ocular formulation to exploit this evolutionary trade-off in clinical settings.

### Circadian clearance modulation enhances time-dependent retention of topical antibiotics

To capture diurnal fluctuations in aqueous humor dynamics, we modeled intraocular drug clearance using a circadian cosine function *ψ*(*t*) calibrated to published aqueous flow data. The resulting waveform, parameterized by a baseline rate, amplitude, and phase shift (Table S1), reproduced the expected oscillatory behavior of aqueous turnover over a 24-hour cycle. The predicted clearance rate was lowest during the early morning hours and increased throughout the day, reaching a maximum near mid-afternoon. These dynamics are consistent with known circadian rhythms in intraocular pressure and aqueous production^41^.

Time-resolved simulations of topical antibiotic elimination revealed a strong dependence of intraocular retention on the phase of administration (Figure 6). For a fixed dose, concentrations decayed more slowly when instillation occurred during periods of reduced clearance, such as the early morning. Conversely, drug applied during high-clearance windows experienced more rapid elimination and reduced exposure duration.

**Figure 6.**
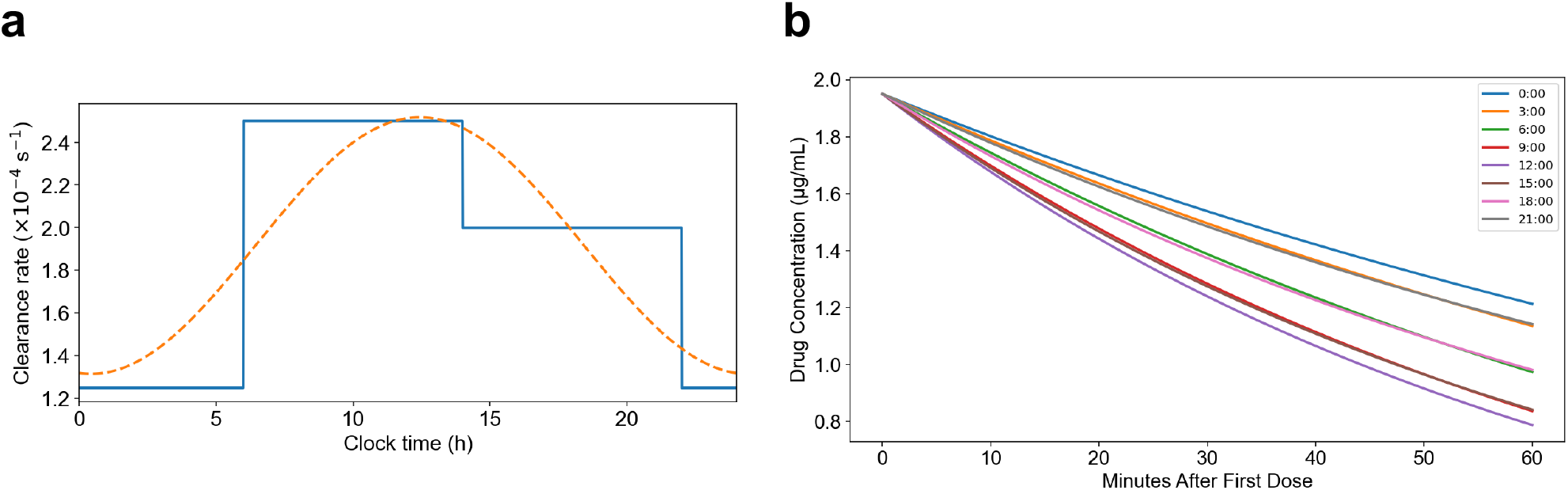
Circadian variation in aqueous humor clearance and its impact on drug pharmacokinetics. **(a)** Circadian modulation of aqueous humor clearance rate (*k*_clear_) modeled as a sinusoidal function (orange dashed line) and approximated by a piecewise step function (blue line) across 24 hours. Clearance rates peak during mid-day and decline during the night, reflecting physiological rhythms in aqueous humor turnover. **(b)** Simulated moxifloxacin drug concentration decay in the aqueous humor over 60 minutes following a single dose administered at different circadian times. Dosing during periods of elevated clearance results in faster drug elimination and lower residual concentrations, while nighttime dosing maintains higher levels due to reduced clearance.

These results indicate that circadian modulation of aqueous turnover significantly alters the pharmacokinetic profile of topical antibiotics. By incorporating this temporal structure into our clearance model, we demonstrate a clear potential for optimizing fluoroquinolone retention through chronotherapeutic scheduling. Aligning prophylactic instillation with lowclearance periods could enhance antimicrobial effectiveness by extending intraocular exposure during early postoperative periods when infection risk is greatest.

### Rotational therapy enhances bacterial suppression under high-resistance conditions

Simulations of moxifloxacin monotherapy every 4 h expose a highly nonuniform drug distribution (Figures S5A–C). Radial profiles peak above 1.5 µg/mL at the corneal surface but drop below 0.5 µg/mL by 6 h at depths >4 mm. Two-dimensional heat maps show drug pooling in the epithelium, leaving mid-aqueous and posterior chambers virtually untreated, while 3D reconstructions reveal persistent posterior “cold spots” that permit bacterial rebound.

By contrast, a four-hour rotation of moxifloxacin and trovafloxacin yields a slightly lower residual vitreous concentration at 24 h (*≈*0.70 µg/mL vs. 0.75 µg/mL) (Figures S6A–F), yet abolishes drug-free intervals: each eight-hour peak fills the other’s trough and maintains anterior levels above the MIC. Although total penetration depth is marginally reduced, the alternating schedule prevents any posterior refuge zones. Limiting peak exposure to a single agent at a time also lowers cumulative drug burden and toxicity, while delivering continuous, spatially uniform inhibitory concentrations that overcome the diffusion limitations of monotherapy.

We compared moxifloxacin monotherapy and rotational fluoroquinolone therapy in simulations to evaluate the impact of treatment strategy on bacterial control under high-resistance conditions (Figures 7A–F). Both regimens administered doses every 4 h, and drug diffusion and bacterial growth were tracked throughout the ocular domain.

**Figure 7.**
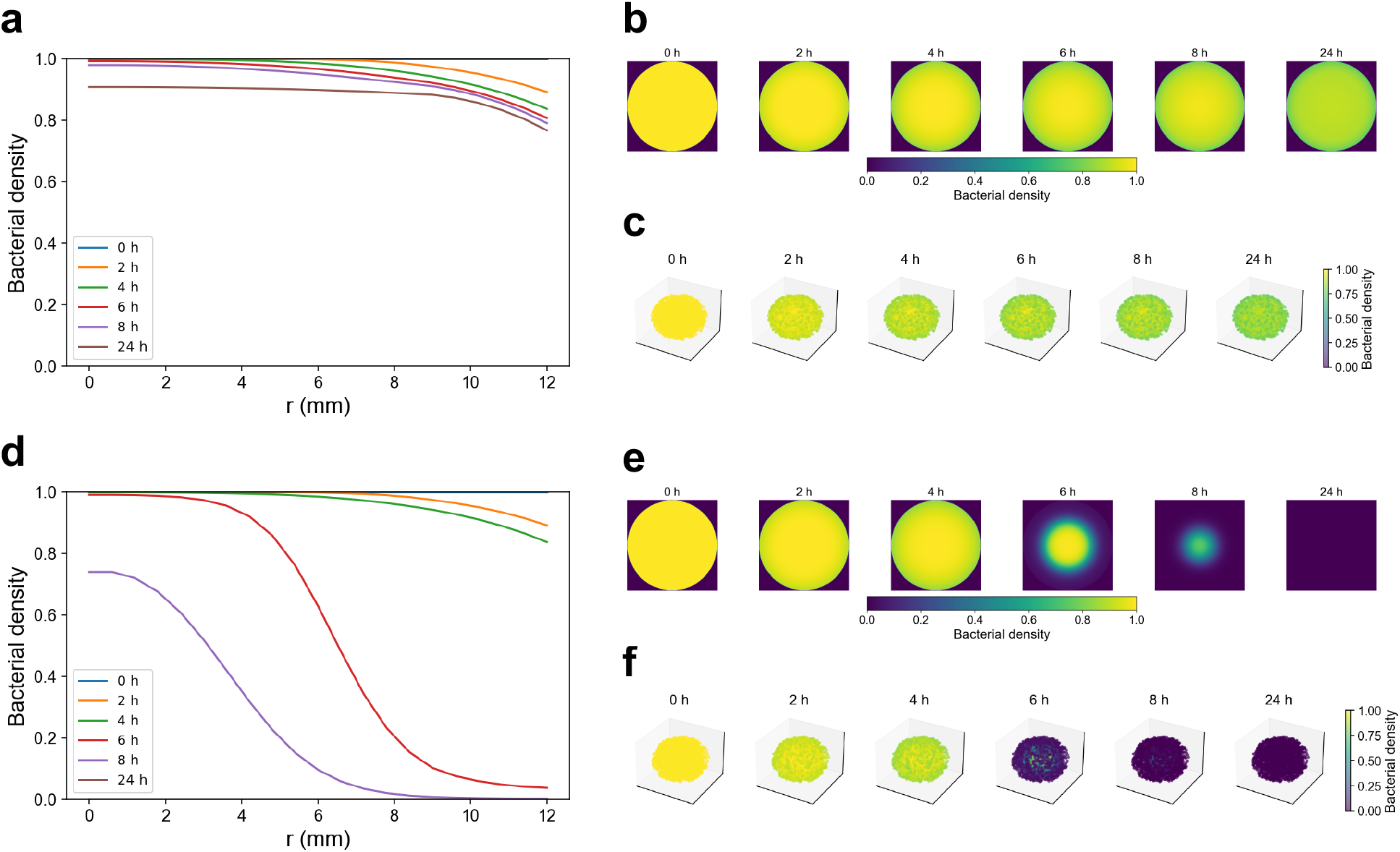
Spatiotemporal dynamics of bacterial suppression under monotherapy and rotational therapy at high resistance with 4-hour dosing intervals. **(a)** Radial bacterial density profiles during monotherapy reveal limited antimicrobial effect, with only slight decreases at the periphery (near *r* = 12 mm) by 24 h and persistent survival in core regions. **(b)** Cross-sectional 2D density maps of the ocular surface confirm marginal suppression over time, with most of the field remaining near baseline density throughout the simulation. **(c)** Corresponding 3D volumetric reconstructions illustrate spatial heterogeneity and lack of deep penetration, with minor bacterial clearance restricted to superficial layers. **(d)** In contrast, rotational therapy produces substantial spatial suppression of the bacterial population, with radial profiles showing a consistent decline in density from surface to core by 24 h. **(e)** 2D cross-sectional maps reveal marked bacterial depletion progressing concentrically from the surface inward, particularly after 24 h of alternating treatment. **(f)** 3D reconstructions demonstrate robust volumetric clearance of bacteria across the entire ocular volume by 24 h.

Under a 4 h dosing regimen of moxifloxacin alone (Figures 7A–C), simulations revealed a brief reduction in bacterial density confined to the corneal surface immediately after each instillation. Radial concentration profiles show sharp peaks at the surface that decay rapidly, and two-dimensional corneal maps confirm minimal penetration into deeper tissues. The three-dimensional reconstruction demonstrates that residual antibiotic is cleared within hours, allowing the bacterial population to rebound fully before the next dose and yielding only slight net decreases in total burden after 24 h.

In contrast, alternating moxifloxacin with trovafloxacin (Figures 7D–F) maintained uninterrupted antimicrobial activity across the ocular domain. Radial peaks of trovafloxacin fill the troughs left by moxifloxacin, while two-dimensional corneal maps illustrate each agent alternately restoring anterior concentrations. The three-dimensional reconstruction confirms that this staggered, dual-drug schedule drives deeper penetration and prevents any interval of bacterial regrowth. Rotational therapy produced a continuous decline in bacterial density and achieved complete eradication by 24 h, overcoming the spatial and temporal limitations of single-agent delivery.

### Spatial gradients and heterogeneity are amplified by rotational therapy

To investigate spatial control dynamics under high-resistance conditions, we compared moxifloxacin monotherapy to rotational fluoroquinolone therapy by examining bacterial distribution and spatial metrics across the ocular domain (Figures 8A-D).

**Figure 8.**
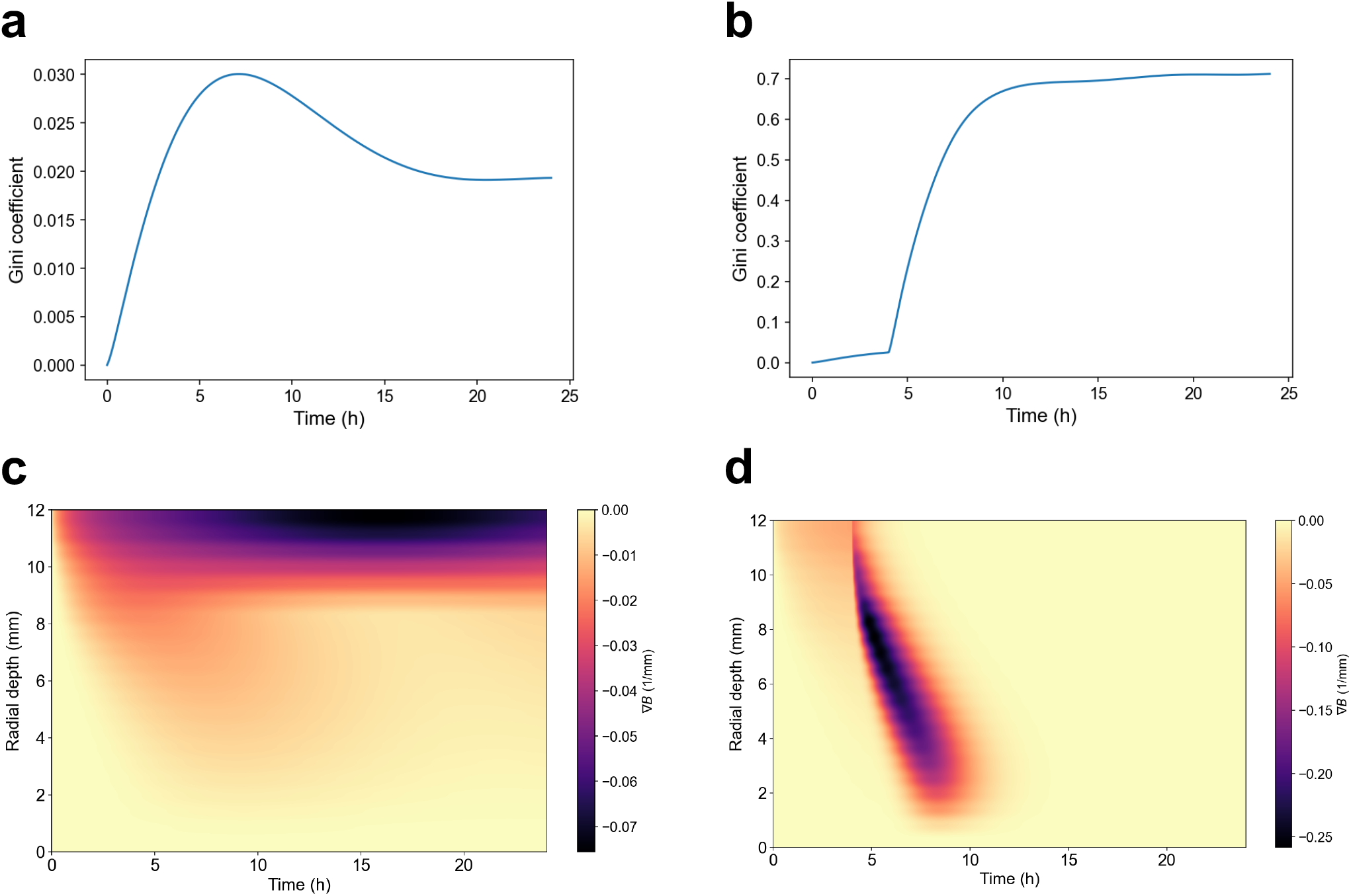
Spatiotemporal heterogeneity in bacterial population dynamics under high-resistance monotherapy and rotational fluoroquinolone treatment. **(a–b)** Global Gini coefficient *G*(*t*) profiles computed across radial bacterial biomass *B*(*r, t*), quantifying total spatial inequality during **(a)** moxifloxacin monotherapy and **(b)** alternating moxifloxacin–trovafloxacin therapy. Higher Gini values indicate more uneven bacterial clearance. **(c–d)** Radial concentration gradients ∇*B* reveal the sharpest biomass depletion zones across space and time for **(c)** monotherapy and **(d)** rotational therapy. Rotational treatment induces rapid and deep bacterial killing, generating steep concentration gradients and fragmented clearance zones early on. By 10 hours, approximately 4 hours after the first pulse of trovafloxacin, the rotational therapy gradient collapses to zero, reflecting complete bacterial clearance. In contrast, monotherapy maintains a small but persistent negative gradient in the outer ocular layers, indicating residual bacterial burden in peripheral regions.

Under monotherapy, bacterial suppression was limited and remained spatially uniform. The Gini coefficient, which quantifies disparity in bacterial density across radial space, stayed low throughout the simulation, reflecting consistently high bacterial load across the ocular tissue. The radial gradient of the bacterial population (∇*B*) remains low and evenly distributed along the full 0 mm to 12 mm depth axis. Each topical dose produces only a brief, shallow increase in the gradient near the corneal surface that quickly returns to baseline before the next instillation. This smooth, undifferentiated profile indicates that drug penetration never sharpens into deeper compartments, which explains why monotherapy fails to achieve complete bacterial eradication.

Additionally, spatial heterogeneity remains minimal across depth and time (Figure S7A–C). The local Gini coefficient stays uniformly low, indicating almost no inequality in bacterial density and a uniformly shallow, incomplete kill. The Laplacian of the bacterial field, ∇^2^*B*, is essentially zero throughout, reflecting the absence of sharply curved “kill fronts” and instead a gradual decay of bacterial density from the corneal surface inward. Likewise, the divergence of bacterial flux,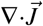, remains near zero at all depths and times, showing that no strong sink regions form where bacterial removal outpaces diffusive back-flow. These results confirm that single-agent monotherapy cannot generate focused, propagating bacterial clearance in any portion of the tissue.

In contrast, rotational therapy induced pronounced spatial heterogeneity. The Gini coefficient increased significantly over time, revealing the emergence of localized bacterial clearance. The gradient of bacterial population, ∇*B*, attains its highest values at the corneal surface and remains elevated through the anterior segment to the vitreous base, reflecting a rapidly advancing kill front; beyond that point bacterial density falls to near zero and ∇*B* likewise becomes zero across the remainder of the ocular domain, demonstrating complete eradication.

The local Gini coefficient displays alternating bands of elevated inequality at shallow depths immediately after each drug switch. Once the bacterial population is eradicated, this coefficient remains uniformly low across all depths. The Laplacian of the bacterial density, ∇^2^*B*, highlights transient regions of strong curvature advancing from the anterior surface into deeper tissues after each dosing interval, which vanish as the density profile flattens. The divergence of the bacterial flux,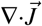, identifies sink regions that coincide with each alternating antibiotic peak and returns to zero once net flux ceases.

These findings demonstrate that rotational therapy does more than uniformly reduce bacterial burden; it creates and propagates steep, localized suppression fronts by alternating spatially distinct kill-zones. By harnessing both temporal offsets and the differing diffusion-clearance profiles of the two fluoroquinolones, this strategy imposes a dynamically heterogeneous environment that may prevent uniform adaptation and resistance development.

## Discussion

The markedly different ocular pharmacodynamics of moxifloxacin and trovafloxacin stem from their distinct structural and physicochemical properties. Moxifloxacin’s 8-methoxy substituent and rigid, bulky C-7 diazabicyclic amine increase electron delocalization and lipophilicity, promoting accumulation in anterior ocular tissues and strong interaction with topoisomerase IV, the primary target in *S. aureus*^23,42^. However, its size and rigidity hinder diffusion into posterior compartments, limiting efficacy under high-resistance conditions.

Trovafloxacin combines a planar 2,4-difluorophenyl moiety at N-1 with a smaller, flexible five-membered nitrogenous ring at C-7. This dual-substituent motif enhances membrane permeation and allows conformational adaptation to varied microenvironments, supporting potent activity even when MICs are elevated in Gram-positive pathogens^17^. Our structural modeling provides mechanistic insight into why moxifloxacin and trovafloxacin respond differently to resistance-conferring mutations: the S80F alteration prevents moxifloxacin’s rigid C-7 diazabicyclic amine from forming hydrophobic contacts, whereas trovafloxacin’s five-membered ring engages Phe80 through multiple hydrophobic and *π − π* stacking interactions, complemented by hydrogen bonds to adjacent residues, thereby preserving high-affinity binding to the mutant enzyme.

Both agents nevertheless carry rare but serious systemic risks. Moxifloxacin can prolong the QT interval via hERG channel blockade^43^ (Figure S10), and trovafloxacin’s systemic use was halted after over 100 reports of idiosyncratic hepatotoxicity, including 14 acute liver failure cases^18,19^. Topical ophthalmic application minimizes systemic exposure, enhancing safety for prophylactic regimens.

A second critical insight from our simulations is the value of chronotherapeutic alignment. Aqueous humor outflow follows a circadian rhythm—with lowest clearance in pre-dawn hours and peak clearance in mid-afternoon—such that administration during low-flow periods prolongs anterior chamber residence time without increasing total daily dose. Synchronizing dosing with periods of maximal bacterial vulnerability may further amplify bactericidal efficacy. Personalized factors such as sleep–wake patterns, glaucoma, inflammation, or epiphora can shift aqueous dynamics and circadian phase. Integrating patient-specific chronobiology, potentially guided by wearable ocular monitors, could refine timing protocols and optimize outcomes.

Despite these promising *in silico* results, several limitations warrant discussion. Our model assumes homogeneous tissue properties and does not account for interpatient variability in corneal permeability or aqueous flow amplitude. We also lack experimental validation of rotational regimens *in vitro* or *in vivo*. A formal sensitivity analysis of key parameters (diffusivities, clearance amplitude, dosing times) is needed to quantify robustness. Finally, clinical translation must consider patient adherence to frequent dosing and potential toxicity of compounded topical formulations.

In conclusion, this study presents a unified computational framework that integrates structural modeling, ligand–protein docking, circadian-informed pharmacokinetic and diffusion simulations, and *in silico* resistance mapping to predict the performance of sequential moxifloxacin and trovafloxacin administration in the anterior segment. Our results indicate that alternating these agents can synergistically exploit their distinct penetration and binding properties, preserve activity against topoisomerase IV mutants, and sustain inhibitory concentrations while reducing systemic exposure. These predictive insights lay the foundation for targeted *in vitro* and *in vivo* investigations of ocular drug distribution and efficacy, and they underscore the value of chronobiologically informed simulations in guiding the design of prophylactic regimens against MRSA.

## Methods

### Baseline Model and Experimental Data Fitting

We use a simple logistic growth model to interpret the experimental bacterial growth data. Denote *B*(*t*) as the bacterial density at time *t* (log CFU/mL):

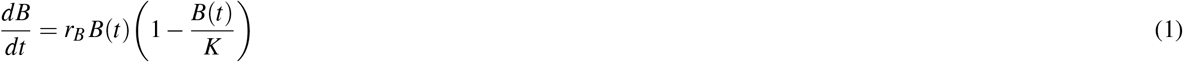

where *r*_*B*_ is the bacterial growth rate (log CFU/mL/hr) and *K* is the carrying capacity (log CFU/mL).

To calculate baseline values and for model calibration purposes, least-square fitting was performed on literature in-vitro MRSA growth rate data^44^. Values were averaged across all isolates tested in the study and fit using equation (1). Using the same literature data as before, Markov Chain Monte Carlo (MCMC) was performed to fit the in-vitro growth rate and obtain posterior distributions.

#### Experimental Data

Experimental bacterial growth data for both untreated and antibiotic-treated conditions were provided by collaborators at the University of Miami Bascom Palmer Eye Institute. Two MRSA isolates were selected based on their response to moxifloxacin exposure: one with a minimum inhibitory concentration (MIC) of 0.25 µg/mL (susceptible) and another with a MIC of 8 µg/mL (resistant). The isolates were cultured and subjected to serial dilution, and bacterial growth was assessed using colony-forming unit (CFU) counts obtained at regular intervals over an 8-hour period. For each timepoint, data were recorded in triplicate, and dilution ranges were selected to ensure countable CFU levels across all measurements.

#### Instrinsic Growth Rate Parameterization

Using the model in equation (1), the experimental growth data without antibiotic treatment for the susceptible strain was plotted in triplicate for each measurement time and fit using least-square fitting. All data points that were uncountable were assumed to be at carrying capacity, which was equal to be 9 log CFU/mL. The growth rate for the susceptible and resistant MRSA strain was then calculated through MCMC by solving differential equation (1).

#### Killing Rate Parameterization

The two MRSA isolates were then re-plated in triplicates onto selective Blood Agar plates with varying concentrations of moxifloxacin. The same procedures as determining the control growth rate were conducted and the isolates underwent the same serial dilutions and CFU assays. Using the estimated growth rate with confidence intervals and posterior distributions as rough initial guesses, MCMC fitting was used on the experimental data. To incorporate the rate of antibiotic killing, the model was updated to:

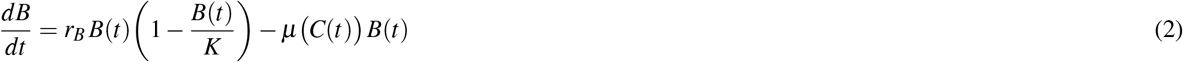

where *µ*(*C*(*t*)) is the antibiotic killing rate at time *t* (log CFU/mL/hr) and *C*(*t*) is the antibiotic concentration at time *t* (g/mL). The following equation was used to calculate and model the antibiotic concentration-dependent killing rate function:

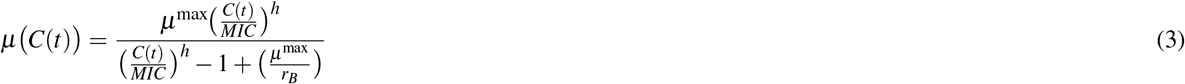

where *µ*^max^ is the maximum antibiotic killing rate (log CFU/mL/hr), *MIC* is the minimum inhibitory concentration (µg/mL), and *h* is the Hill coefficient.

We assume that the antibiotic concentration remains constant during the kill curve experiments; that is, *C*(*t*) is treated as a time-independent constant for the purpose of data fitting. Kill-curve graphs were produced for each moxifloxacin concentration tested (0.125, 0.5, 2, 8, and 32 µg/mL) and the killing rate was calculated using MCMC fitting for each condition. Then, moxifloxacin killing rate was plotted as a function of antibiotic concentration. MCMC posterior distributions were calculated in the same manner to fit and estimate unknown parameters in equation (3).

### Topoisomerase IV binding

#### Docking Simulations

In order to simulate and analyze our proposed mechanism of collateral sensitivity, ligand-target docking simulations were conducted. The three-dimensional structure of MRSA topoisomerase IV (wild type) was downloaded from the Protein Data Bank.

To model fluoroquinolone resistance, the wild-type topoisomerase IV subunit A (2INR)^45^ and subunit B structures (4URL)^46^ was imported into PyMOL^47^. The target residue corresponding to serine at position 80 (a mutation hotspot in the QRDR of ParC) was identified and mutated to phenylalanine (S80F), generating a model of a resistant topoisomerase IV^33,48,49^. The mutated structure was then imported into UCSF Chimera for energy minimization^50^. A brief minimization was performed to relieve any unfavorable steric interactions introduced by the mutation while maintaining the overall protein conformation. The minimized model was saved and subsequently used for docking simulations. Both the wild-type and the S80F-mutated topoisomerase IV models were submitted to SwissDock for comparative docking studies^51,52^. Identical grid settings and scoring parameters were used to dock moxifloxacin and trovafloxacin into the binding pocket of each enzyme variant. The docking results, including predicted binding affinities and ligand orientations, were then analyzed to assess how the S80F mutation alters drug binding and to explore the potential for collateral sensitivity.

#### Statistical Analysis

All statistical analyses were performed using GraphPad Prism 10 and in accordance with standard reporting guidelines. Binding energy data from four experimental groups were analyzed using one-way analysis of variance (ANOVA) to assess differences in group means. ANOVA was selected due to its suitability for comparing more than two groups under the assumption of normally distributed data with equal variances.

To justify use of parametric testing, we verified the assumption of homogeneity of variances using both the Brown–Forsythe and Bartlett’s tests. Normality of residuals was assessed by visual inspection of Q–Q plots and histograms. All tests were two-tailed, and the significance threshold (*α*) was set at 0.05.

Where ANOVA indicated statistically relevant group-level differences, post hoc comparisons were conducted using Tukey’s honestly significant difference (HSD) test to correct for multiple comparisons. This approach was chosen to control the family-wise Type I error rate while comparing all pairwise group combinations.

Descriptive statistics are reported for each group as mean ± standard deviation (s.d.), with sample sizes (*n*) explicitly stated in figure legends and results. Graphs include error bars that reflect s.d., not standard error of the mean, unless otherwise noted.

### Circadian modeling of aqueous humor clearance

To model the time-dependent clearance of topically administered drugs from the anterior chamber, we constructed a smooth circadian clearance function based on literature-reported aqueous humor flow rates^53^. The anterior chamber was assumed to have a fixed volume of *V*_AC_ = 200 *µ*L. The flow rate *F*(*t*) (in *µ*L/min) varies over a 24-hour cycle to reflect circadian modulation in aqueous production. These time-dependent flow rates were converted into first-order clearance rate constants *k*(*t*) (in s^*−*1^) using the relationship:

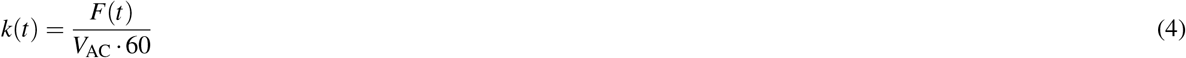

yielding a piecewise function *k*(*t*) (in s^*−*1^) sampled every minute over a 24-hour period. To facilitate analytical integration in downstream pharmacokinetic models, we fit this profile with a smooth circadian waveform:

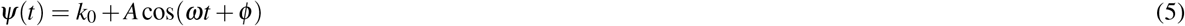

where *k*_0_ is the baseline clearance, *A* is the modulation amplitude, 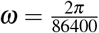 is the circadian frequency (rad/s), and *ϕ* is the phase shift.

To obtain *A* and *ϕ*, we first performed least-squares regression using a first-order Fourier approximation:

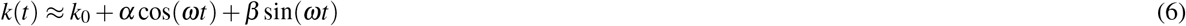

where *α* and *β* are regression coefficients with units of s^*−*1^, capturing the cosine and sine components of circadian variation.

The fitted coefficients were then transformed into physiologically interpretable amplitude and phase shift values:

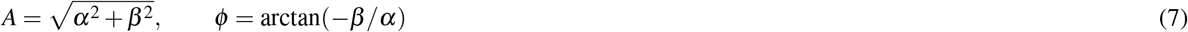

To quantify the impact of dosing time on intraocular drug retention, we simulated drug concentration decay in the anterior chamber following a single topical dose. For each simulated dosing time across the 24-hour cycle, drug elimination was computed by solving the first-order decay equation:

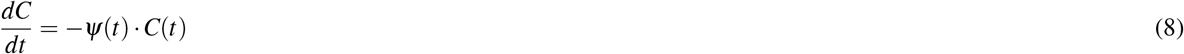

The initial condition *C*(0) was set to a fixed bolus concentration, and the equation was integrated numerically. This approach enabled direct comparison of exposure profiles for identical doses administered at different circadian phases, capturing the influence of biologically driven variations in aqueous humor turnover on early drug availability.

### Spatiotemporal modeling of drug diffusion and bacterial killing

We modeled the anterior segment of the eye as a radially symmetric spherical domain of radius *R* = 12 mm, representing concentric layers of the cornea, aqueous humor, and vitreous humor. Let *r∈*[0, *R*] denote the radial coordinate and *C*_*X*_ (*r, t*) the concentration of a topically administered fluoroquinolone *X ∈ {A, B}*, where *A* represents moxifloxacin and *B* represents trovafloxacin. Each drug was simulated using a radial diffusion equation with spatially heterogeneous diffusivity and aqueous humor clearance:

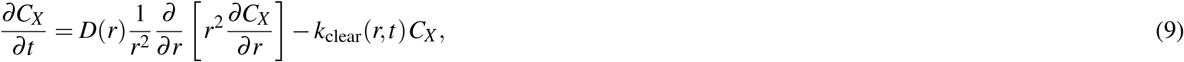

where *D*(*r*) reflects tissue-specific diffusivities and *k*_clear_(*r, t*) denotes first-order clearance localized to the aqueous humor. We assume a homogeneous clearance rate across the eye and use the function *ψ*(*t*), as parametrized in the previous section, for the simulation.The function *ψ*(*t*) was applied to clearance rates within the aqueous layer, while corneal and vitreous compartments were assigned constant clearance of zero.

We adopt the following Neumann boundary and initial conditions under the assumptions that (i) there is no drug loss through at the tear film–air interface located in the anterior surface of the cornea, (ii) there is zero flux at the geometric center of the vitreous body, and (iii) antibiotics are uniformly applied over tear-air interface at the onset of treatment:

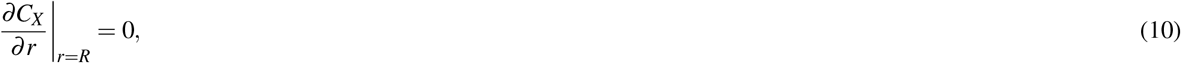

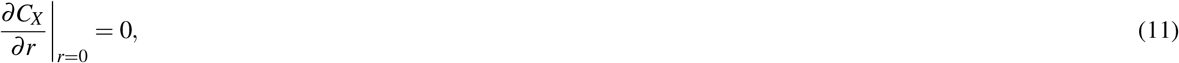

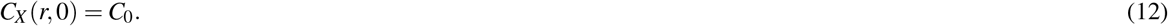

In monotherapy, only drug A was instilled every 4 hours. In rotational therapy, A and B alternated every 4 hours, starting with drug A and continuing cyclically.

The bacterial population *B*(*r, t*) followed logistic growth with drug-induced killing:

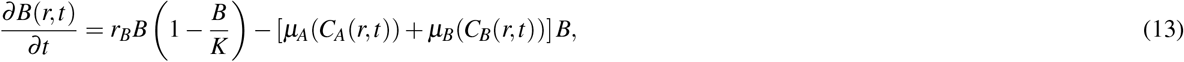

where *r*_*B*_ is the baseline growth rate, *K* is the nondimensional carrying capacity, and *µ*_*X*_ (*C*) is the kill function of drug *X* defined by a Hill-type sigmoidal model:

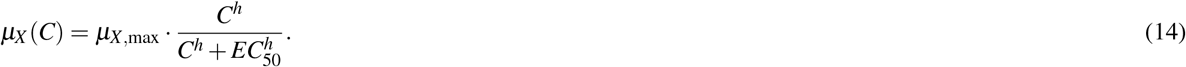

The primary efficacy outcome was total bacterial burden over time, computed as:

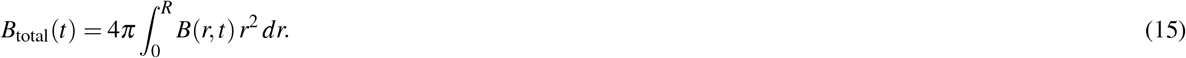

### Spatial heterogeneity analysis

To quantify spatial heterogeneity in bacterial clearance, we computed the global Gini coefficient *G*(*t*) at each timepoint *t*, defined as:

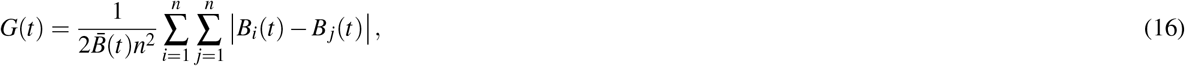

where *B*_*i*_(*t*) is the bacterial density at radial node *r*_*i*_, 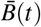 is the mean bacterial load across all *n* spatial positions at time *t*, and the double summation captures all pairwise differences in bacterial abundance. Higher values of *G*(*t*) indicate greater spatial disparity in bacterial suppression, arising from nonuniform drug exposure.

We also analyzed local Gini coefficients *G*_local_(*r*_*i*_, *t*) to assess heterogeneity within subregions of the anterior chamber. This was computed using a sliding window centered at each spatial node *r*_*i*_, incorporating its immediate neighbors (e.g., a 5-point stencil) to estimate localized inequality in bacterial burden:

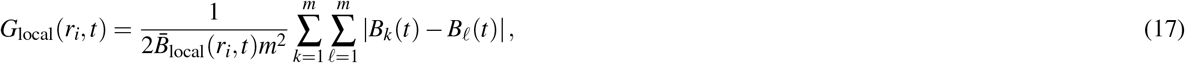

where *m* is the number of nodes in the local window and 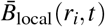 is the mean bacterial density in that neighborhood. By mapping *G*_local_ over space and time, we captured the formation of spatial hotspots and depletion zones during treatment.

To further explore spatial dynamics, we computed the radial gradient of bacterial density:

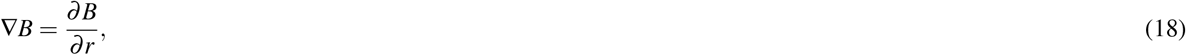

which captures the direction and steepness of changes in bacterial population across radial space. Regions with steeper gradients indicate sharp bacterial clearance fronts.

We also calculated the Laplacian to assess curvature of the bacterial profile:

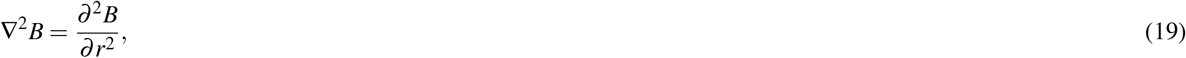

identifying convex and concave patterns in spatial bacterial suppression.

Fick’s First Law defines the bacterial flux 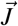 due to diffusion as:

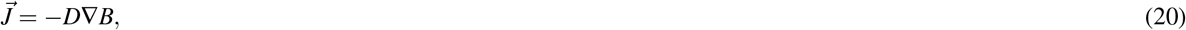

where *D* is the diffusion coefficient. From this, we compute the divergence of flux:

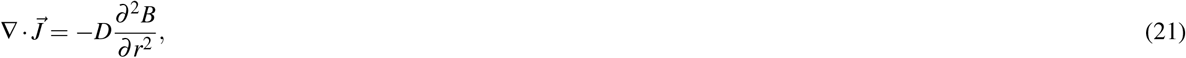

which reveals spatiotemporal regions where bacterial density is actively depleted (sink) or accumulated (source) by diffusion.

## Acknowledgements

This research was partially supported by the National Science Foundation DMS-2052648. Additionally, we acknowledge the use of BioRender and ChemDraw in the preparation of select figures included in this manuscript.

## Data availability statement

The complete simulation files and raw docking data are publicly available at: https://doi.org/10.5281/zenodo.15694462.

## Author contributions statement

A.S. designed and conducted the simulations and wrote the original manuscript draft. D.M. curated and compiled the experimental data. X.H. assisted in the simulation process and edited the manuscript.

## Notes

https://doi.org/10.5281/zenodo.15694462.

### Competing Interest Statement

The authors have declared no competing interest.

