## Supplemental Information for "Modeling Rotational Fluoroquinolone Therapy as a Novel Treatment for Ophthalmic MRSA Infections"

### Supplementary Information

#### MCMC Fitting

We used the Python `emcee` package with an ensemble of 100 walkers sampling the growth-rate model's two parameters, $r \sim U(0,1)$ and $\sigma \sim U(0,3.6)$, initialized near $(0.5, 0.6)$, for 1,000 steps (after a brief burn-in), and similarly ran 50 walkers for 1,000 steps to infer the killing-rate parameters $(\mu, k)$ under uniform priors (Fig. S3 - S4).

#### Trovafloxacin $C_{max}$ Estimation

To estimate the peak aqueous concentration of trovafloxacin, we applied the literature-reported aqueous/plasma $\mathrm{C_{max}}$ ratio for moxifloxacin in humans (44.3 %) [2]. Thus:

$$C_{\mathrm{max,aq}} = C_{\mathrm{max,plasma}} \times \frac{C_{\mathrm{max,aq}}}{C_{\mathrm{max,plasma}}} = 2.3\ \mu\mathrm{g/mL} \times 0.443 \approx 1.02\ \mu\mathrm{g/mL}.$$

We therefore use $C_{\mathrm{max,aq}} = 1.02\ \mu$g/mL for our spatiotemporal PK/PD models of topical trovafloxacin (Table S1).

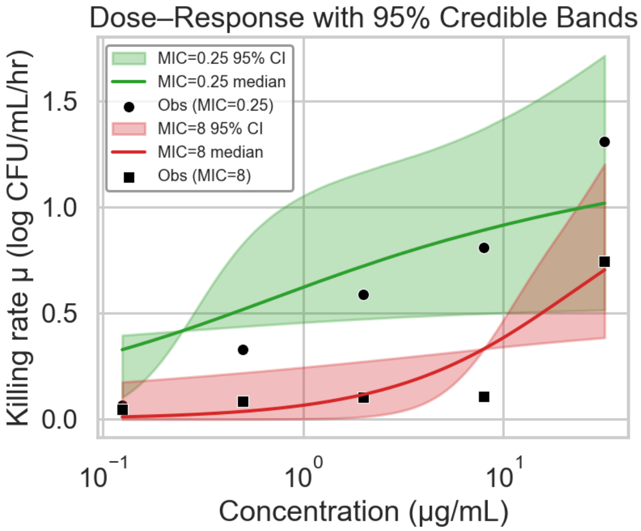


Figure S1: **Posterior dose-response relationships for moxifloxacin-induced killing of MRSA.** Solid green and red lines show the posterior median killing rate $\mu(A)$ for MIC=0.25µg/mL and MIC=8µg/mL, respectively; the corresponding shaded bands are the pointwise 95% credible intervals. Black circles and squares are the observed killing rates at each concentration for MIC=0.25 and MIC=8, respectively. The susceptible isolate (MIC=0.25) exhibits a steep increase in $\mu$ just above its MIC, whereas the resistant isolate (MIC=8) shows a much flatter response and requires higher drug levels to approach its maximal killing rate.

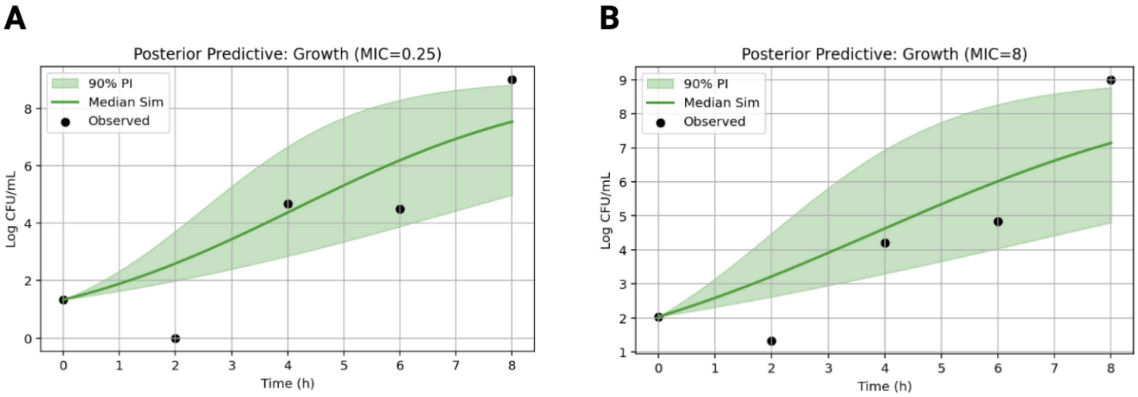


Figure S2: **Posterior predictive checks for MRSA growth under moxifloxacin-free conditions.** Shaded bands show the pointwise 90% predictive intervals, solid lines the median simulation, and black dots the observed log CFU/mL data. Panel (**A**) is for the susceptible isolate (MIC = 0.25µg/mL) and panel (**B**) for the resistant isolate (MIC = 8 µg/mL).

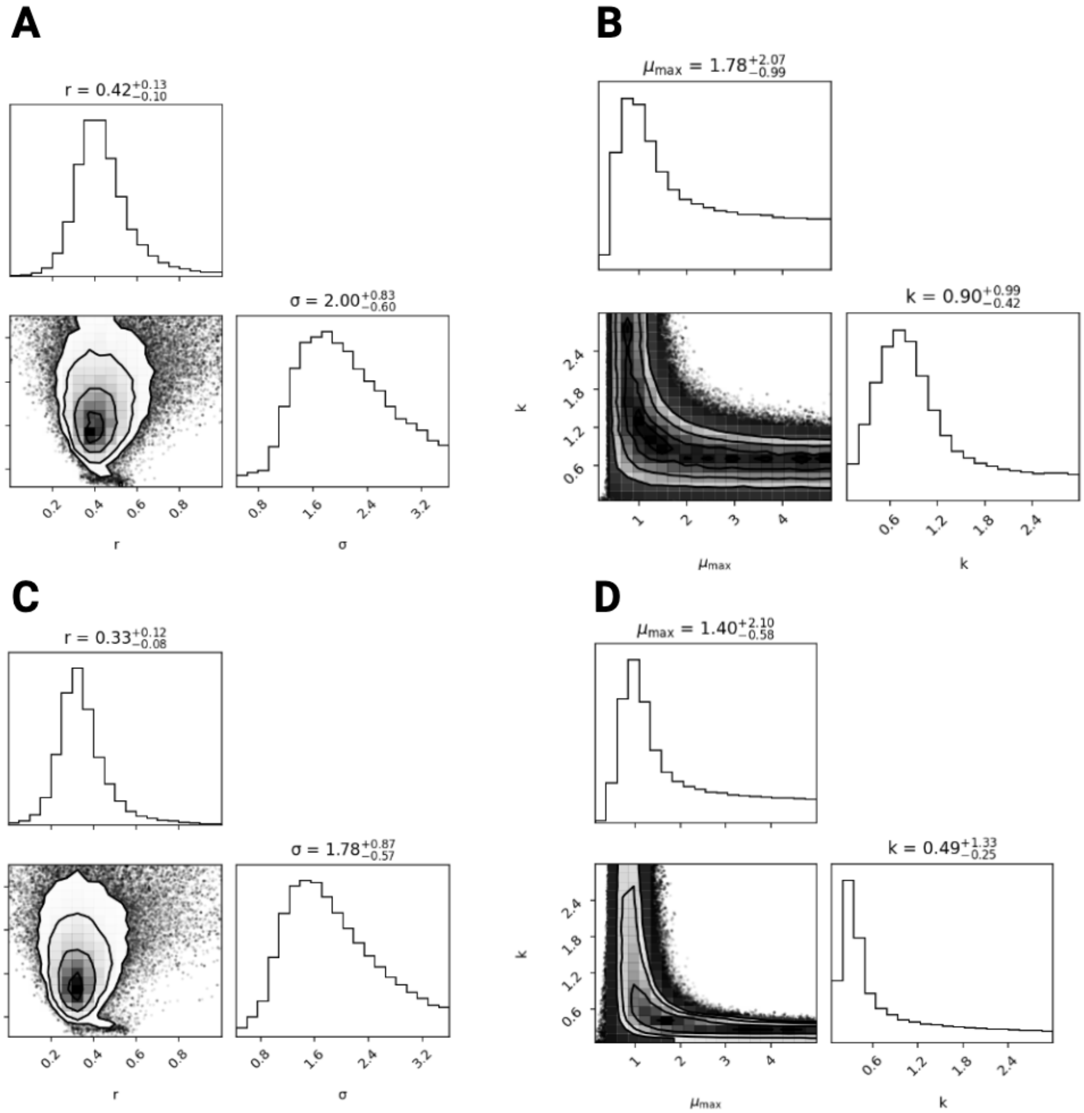


Figure S3: **Marginal and joint posterior distributions of MCMC-estimated parameters for MRSA growth and killing.** (**A**) Susceptible isolate (MIC = 0.25 µg/mL) growth: top panel shows the marginal posterior of the growth rate $r$; lower left shows the joint posterior of $r$ and noise level $\sigma$; lower right shows the marginal posterior of $\sigma$. (**B**) Susceptible isolate (MIC = 0.25 µg/mL) kill: top panel shows the marginal posterior of the maximum kill rate $\mu_{\max}$; lower left shows the joint posterior of $\mu_{\max}$ and Hill coefficient $k$; lower right shows the marginal posterior of $k$. (**C**) Resistant isolate (MIC = 8 µg/mL) growth: top panel shows the marginal posterior of $r$; lower left shows the joint posterior of $r$ and $\sigma$; lower right shows the marginal posterior of $\sigma$. (**D**) Resistant isolate (MIC = 8 µg/mL) kill: top panel shows the marginal posterior of $\mu_{\max}$; lower left shows the joint posterior of $\mu_{\max}$ and $k$; lower right shows the marginal posterior of $k$.

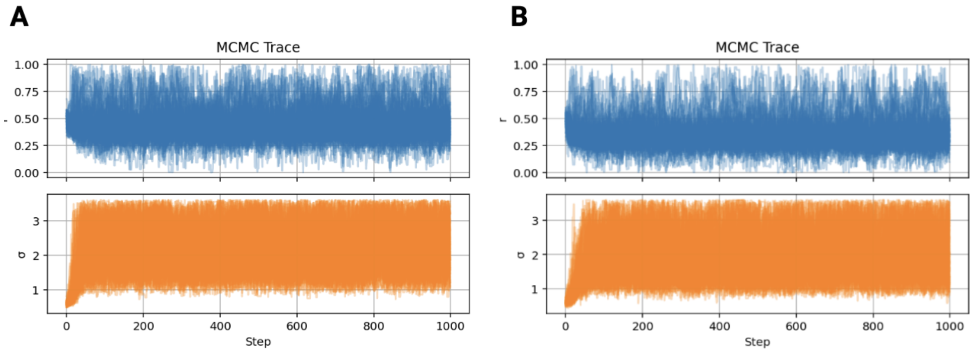


Figure S4: **MCMC trace plots for kill-slope parameter estimation.** Panel (**A**) shows the ensemble sampling traces for the susceptible isolate (MIC = 0.25 μg/mL): the upper trace is the growth rate $r$ and the lower trace is the noise parameter $\sigma$. Panel (**B**) shows the corresponding traces for the resistant isolate (MIC = 8 μg/mL). Both panels illustrate good mixing and stationarity over 1,000 MCMC steps.

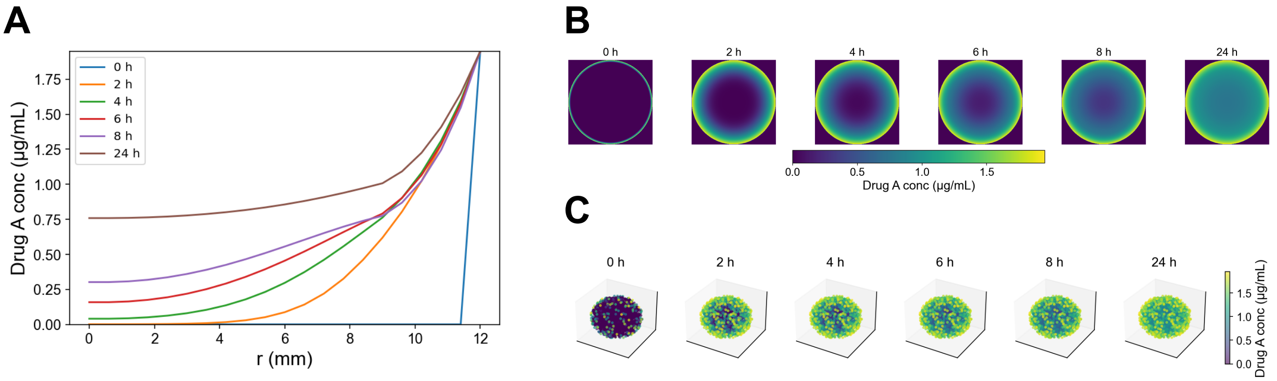


Figure S5: **Spatiotemporal distribution of moxifloxacin in the ocular domain under monotherapy with 4-hour dosing intervals for a high-resistance strain**. (**A**) Radial concentration profiles at 0, 2, 4, 6, 8, and 24h post-first dose, showing steep anterior gradients and rapid decline over time. (**B**) 2D cross-sectional heat-maps of corneal surface concentration, illustrating reduced posterior penetration. (**C**) 3D volumetric reconstructions of intraocular drug distribution, highlighting reduced delivery to deeper compartments and increased aqueous clearance.

Table S1: Model parameters for drug diffusion, clearance, and bacterial dynamics.

| Symbol | Value | Units | Description | Reference |
|---|---|---|---|---|
| *Geometry and Diffusion* | | | | |
| $R$ | 12 | mm | Total eye radius | [1] |
| $\delta_c$ | 0.55 | mm | Corneal thickness | [1] |
| $\delta_a$ | 2.53 | mm | Aqueous humor thickness | [12] |
| $D_{\text{cornea}}$ | $5.84 \times 10^{-10}$ | $\text{m}^2/\text{s}$ | Diffusivity in cornea | [8] |
| $D_{\text{aqueous}}$ | $8.23 \times 10^{-10}$ | $\text{m}^2/\text{s}$ | Diffusivity in aqueous humor | [9] |
| $D_{\text{vitreous}}$ | $3.4 \times 10^{-10}$ | $\text{m}^2/\text{s}$ | Diffusivity in vitreous | [7] |
| $k_{\text{aq}}$ | $2.35 \times 10^{-4}$ | $\text{s}^{-1}$ | Baseline aqueous clearance | [4] |
| *Circadian Clearance Modulation* | | | | |
| $k_0$ | $1.917 \times 10^{-4}$ | $\text{s}^{-1}$ | Baseline clearance | Modeled |
| $A$ | $6.008 \times 10^{-5}$ | $\text{s}^{-1}$ | Amplitude | Modeled |
| $\phi$ | 12.43 | hr after midnight | Phase shift | Modeled |
| *Initial and Boundary Conditions* | | | | |
| $C_X(r, 0)$ | 0 | µg/mL | Initial drug concentration | Assumed |
| $B(r, 0)$ | 1.0 | unitless | Initial bacterial density | Assumed |
| $\tau_A$ | every 4 hr | hr | Dosing interval for moxifloxacin | Modeled |
| $\tau_T$ | every 4 hr (alternating) | hr | Dosing interval for trovafloxacin | Modeled |
| *Trovafloxacin Pharmacodynamics* | | | | |
| $\mu_{\max}$ | 1.85 | log CFU/mL/hr | Max. killing rate | [10] |
| $C_{\max,T}^{\text{plasma}}$ | 2.3 | µg/mL | Max. plasma conc. | [11] |
| $C_{\max,T}^{\text{aqueous}}$ | 1.02 | µg/mL | Estimated aqueous conc. | Calculated |
| $h_T$ | 1.1 | unitless | Hill coefficient | Assumed |
| $EC_{50,T}$ | 0.06 | µg/mL | Half-max. effective conc. | [3] |
| *Moxifloxacin Pharmacodynamics* | | | | |
| $C_{\max,A}$ | 1.95 | µg/mL | Max. aqueous conc. | [5] |
| $EC_{50,A}$ | $6.76 \times$ MIC | µg/mL | Half-max. effective conc. | [6] |

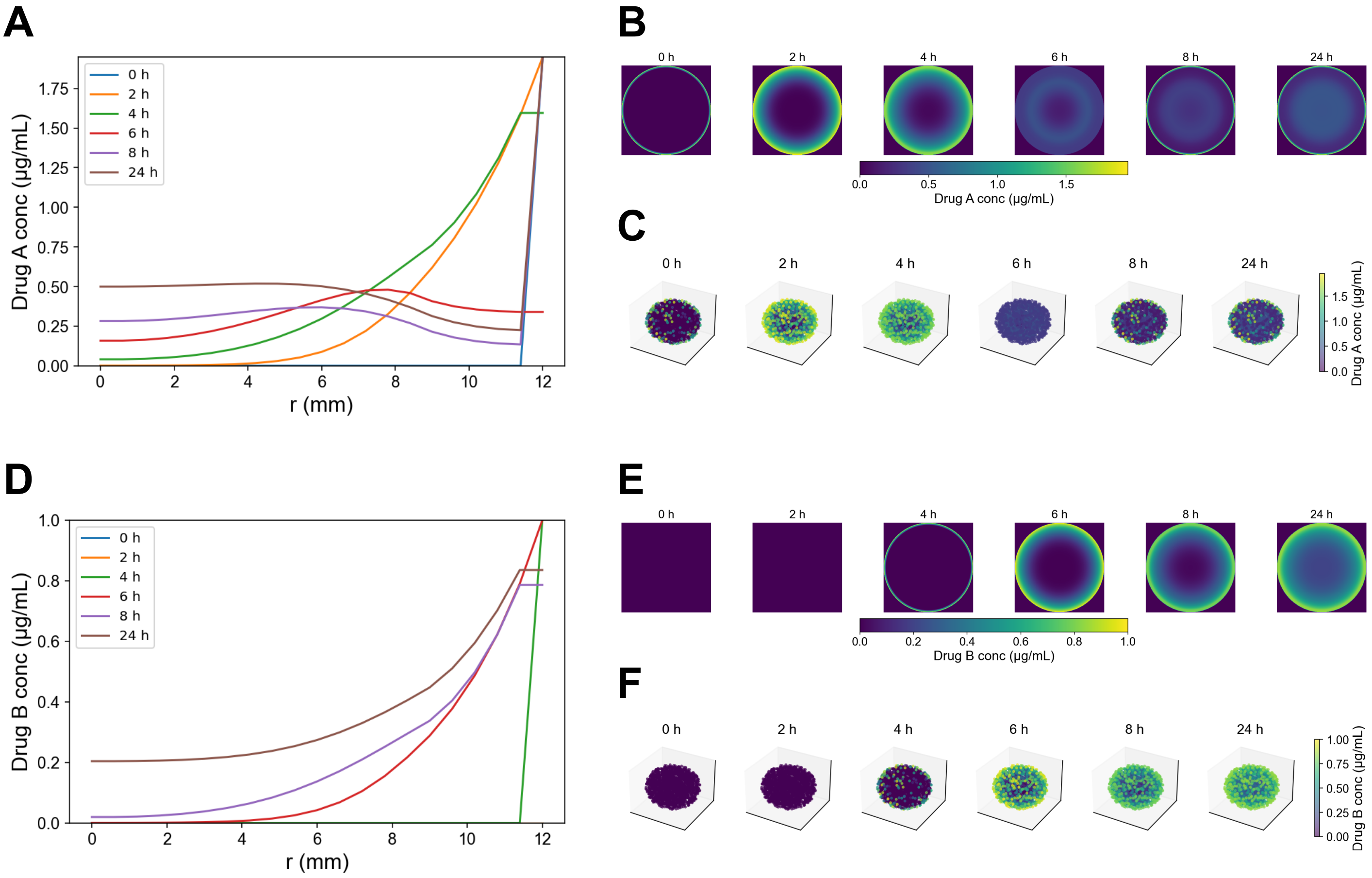


Figure S6: **Spatiotemporal distribution of Drug A (moxifloxacin) and Drug B (trovafloxacin) in the rotational regimen under high-resistance conditions with 4h dosing intervals. Drug A (moxifloxacin) profiles**: (**A**) Radial concentration of Drug A at 0, 2, 4, 6, 8, and 24h, showing its initial peaks and subsequent decay; (**B**) 2D corneal heat-maps of Drug A, illustrating its anterior distribution and limited posterior penetration; (**C**) 3D volume renderings of Drug A, highlighting spatial confinement to anterior compartments. **Drug B (trovafloxacin) profiles**: (**D**) Radial concentration of Drug B at 0, 2, 4, 6, 8, and 24h, showing complementary peaks that overlap Drug A's troughs; (**E**) 2D corneal heat-maps of Drug B, demonstrating sequential anterior coverage by the second agent; (**F**) 3D volume renderings of Drug B, depicting reduced distribution into deeper regions.

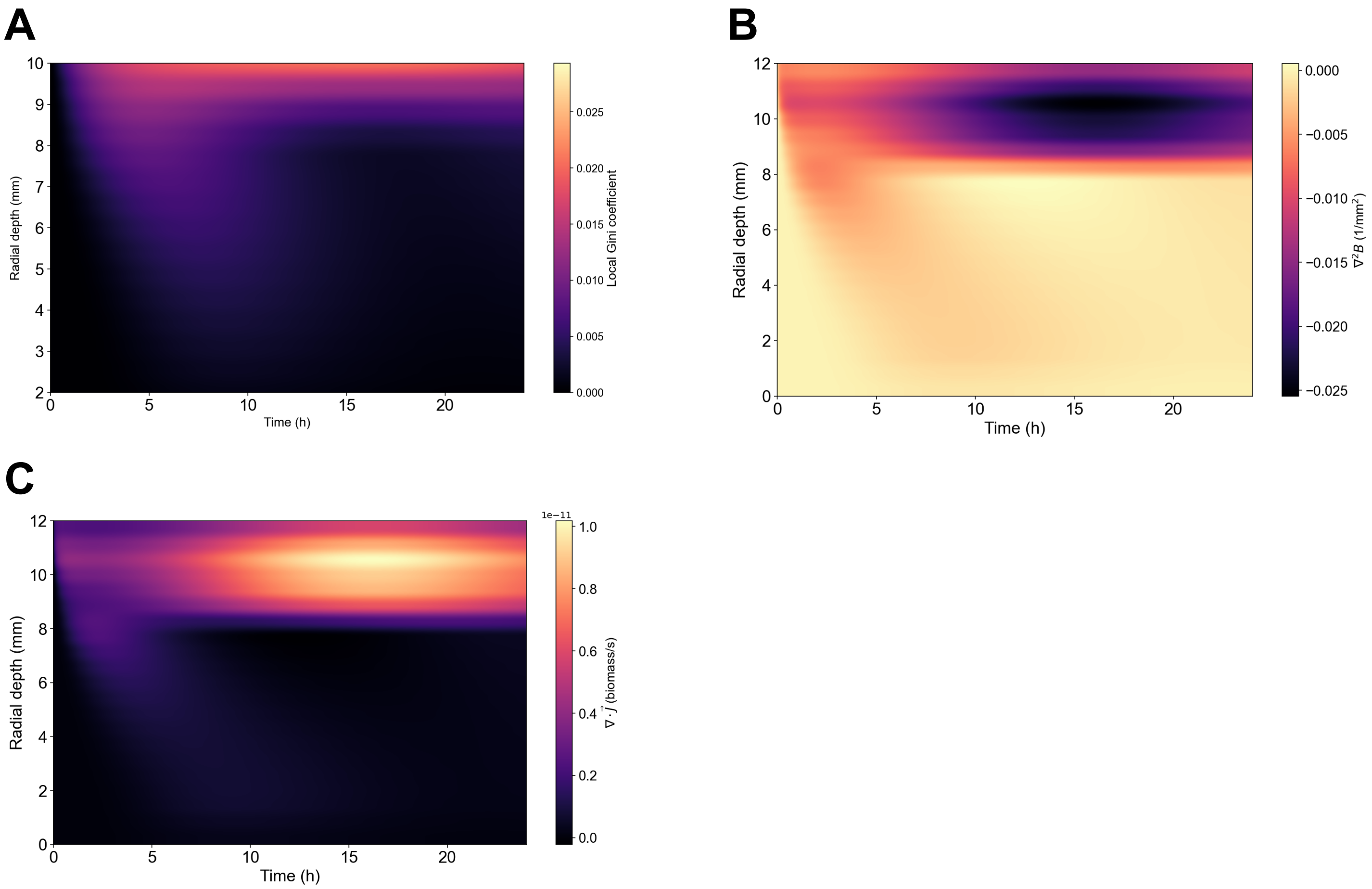


Figure S7: **Spatial heterogeneity metrics under moxifloxacin monotherapy (4h intervals) for a high-resistance strain**. (**A**) Local Gini coefficient over time and radial depth, quantifying inequality in bacterial density and revealing uniformly low heterogeneity under single-agent dosing. (**B**) The Laplacian, $\nabla^2 B$, measures the curvature of bacterial suppression fronts; its near-zero values at increased depths indicate a gradual decay profile rather than the presence of sharply defined clearance boundaries. (**C**) Divergence of bacterial flux ($\nabla \cdot \vec{J}$), highlighting minimal sink regions and showing that diffusion alone cannot overcome sub-MIC exposure zones under monotherapy.

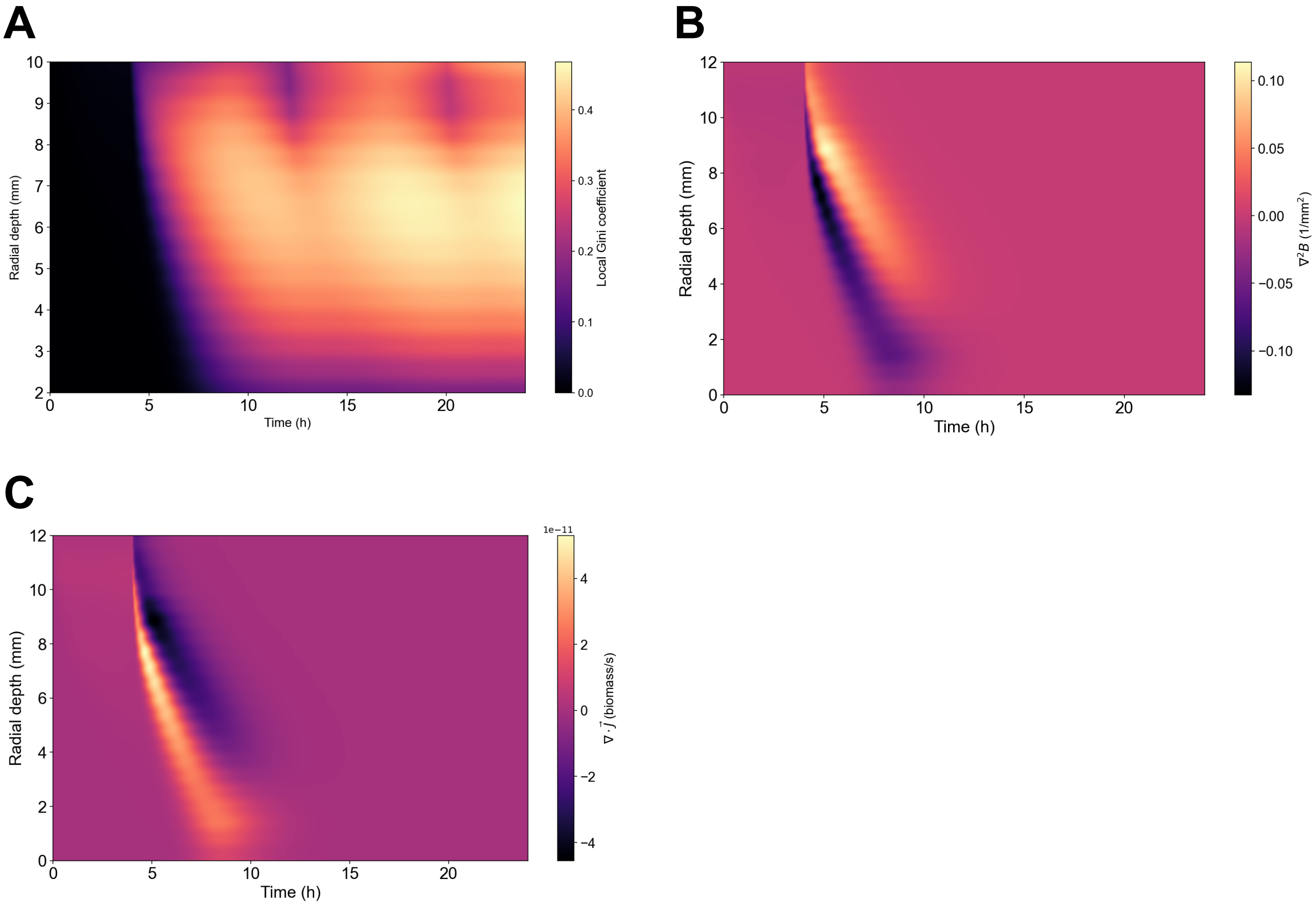


Figure S8: **Spatial heterogeneity metrics under rotational fluoroquinolone therapy (4h intervals) for a high-resistance strain**. (**A**) Local Gini coefficient over time and radial depth, showing alternating bands of elevated inequality that emerge after each 4h drug switch and penetrate from the corneal surface into the vitreous before subsiding once bacterial density falls to zero. (**B**) Laplacian of the bacterial density field, revealing a pronounced, finger-shaped region of high curvature that advances from the anterior cornea to approximately 10mm depth during each suppression front and then returns to near zero as the tissue is sterilized. (**C**) Divergence of bacterial flux ($\nabla \cdot \vec{J}$), identifying transient sink regions aligned with each alternating drug peak, which collapse to zero once bacterial eradication is complete.

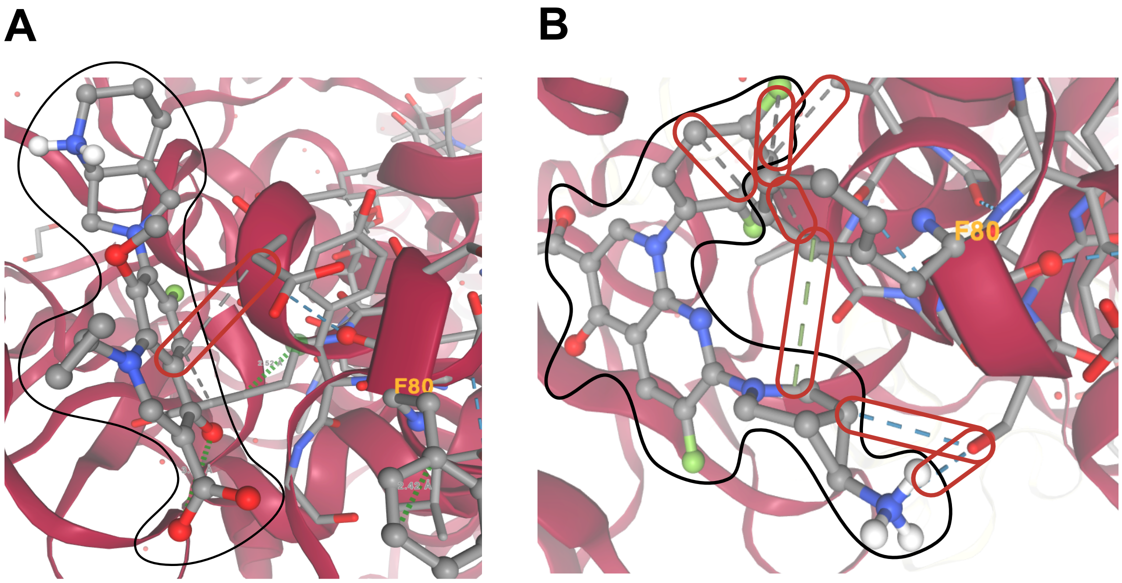


Figure S9: **Differential docking of fluoroquinolones to S80F-mutant topoisomerase IV.** (**A**) Moxifloxacin exhibits only a single hydrophobic contact with a non-mutated residue in the binding pocket and is unable to accommodate Phe80, resulting in poor binding affinity. (**B**) Trovafloxacin's five-membered nitrogenous ring and planar 2,4-difluorophenyl substituent engage Phe80 directly through multiple hydrophobic and $\pi - \pi$ stacking interactions (dashed gray and green lines), while additional hydrogen bonds to adjacent residues (blue dashed lines) further stabilize its binding to the mutant enzyme.

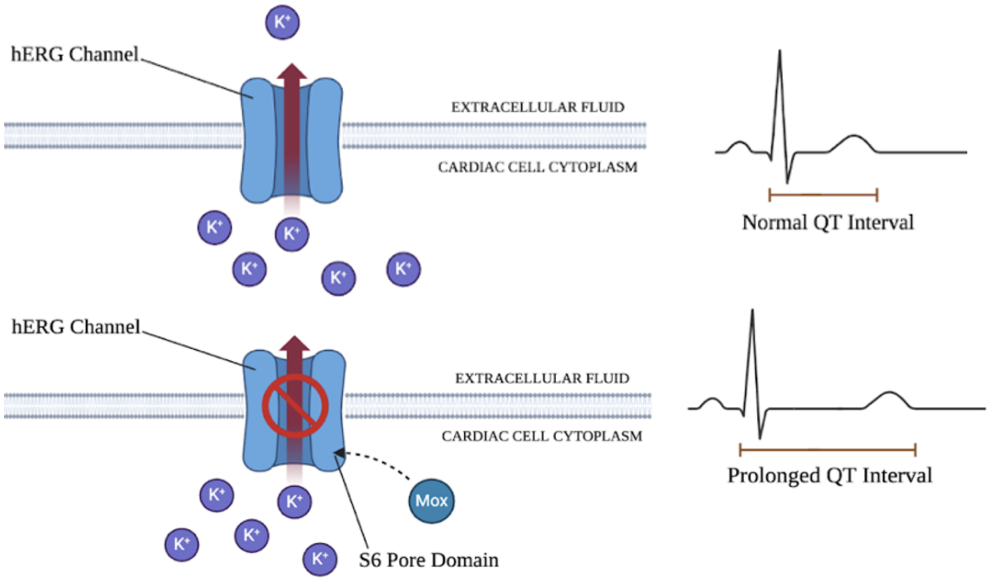


Figure S10: **Moxifloxacin-induced QT interval prolongation**. **(Top)** Normally functioning cardiac hERG channel which undergoes adequate repolarization of potassium ions. **(Bottom)** Upon binding of moxifloxacin at the S6 pore domain of the cardiac hERG channel, an inhibitory effect is seen. The repolarization time is significantly extended and results in a prolonged QT interval.
